# Articular calcified cartilage microarchitecture has a predictive role in abnormal osteochondral strain distributions in murine age-related osteoarthritis

**DOI:** 10.64898/2026.09.18.752585

**Authors:** Lucinda AE Evans, Aikta Sharma, Jishizhan Chen, Lucie E Bourne, Alissa L Parmenter, Joseph Brunet, Kamel Madi, Sebastian Marussi, Peter D Lee, Andrew A Pitsillides, Katherine A Staines

**Author notes:** Corresponding authors (PDL, AAP, KAS), PDL -; AP -; KS. **Email addresses:** LAEE -; AS –; JC -; LEB –; ALP -; JB -: KM -; SM. **Author contributions**. Conceptualisation: LAEE, AS, PDL, AAP, KAS; Methodology: LAEE, AS, LEB, KM, JC, ALP, PDL, AAP, KAS; Software: LAEE, AS, LEB, KM, JC, JB; Validation: LAEE, AS; Formal analysis: LAEE, AS, LEB, JC; Investigation: LAEE, AS, LEB, JC, ALP, JB; Resources: PDL, AAP and KAS; Data curation: LAEE, AS; Writing – original draft: LAEE, AS, KAS; Writing – review and editing: all authors; Visualisation: LAEE, AS, JC; Supervision: PDL, AAP and KAS; Project administration: PDL, AAP and KAS; Funding acquisition: PDL, AAP and KAS.

## Abstract

Even though osteoarthritis is considered a disease of the whole joint, integrated analysis of articular calcified cartilage (ACC) behaviour with adjacent tissues remains a persistent limitation. Herein, we performed synchrotron X-ray computed tomography in intact loaded joints and coupled anatomical analysis with digital volume correlation to examine whether microarchitectural ACC features serve as predictive mechano-biomarkers of osteoarthritis. We reveal that male osteoarthritis-prone STR/Ort mice have thicker yet more porous ACC that contains larger but less spherical chondrocyte lacunae than their healthy parental-control CBA mice. This was linked to asymmetrical distribution of tensile and compressive strains generated under physiological loads in the ACC and underlying subchondral bone in tibial epiphyses of STR/Ort mice. Together these data suggest that murine ACC microarchitecture has a mechano-predictive role in age-related osteoarthritis.

## Introduction

Primary osteoarthritis (OA) is a profoundly debilitating, age-related disease, with rising prevalence contributing substantially to the global burden of musculoskeletal disability.^1,2^ Despite extensive research, the precise anatomical and mechanical factors, which initiate and drive OA in humans and in animals remain unclear.^3^ In recent years, attention has shifted from the articular cartilage to other neighbouring joint tissues including the subchondral bone (SCB), synovium and ligaments as the field adopts a more unified understanding of OA as a disease of the ‘whole joint’.^4^

However, the articular calcified cartilage (ACC) - a layer deep to the hyaline articular cartilage (HAC) and superficial to the SCB - is often overlooked.^5,6^ Research however has highlighted that morphologic, metabolic and biomechanical ACC features may play in important role in OA pathophysiology.^7–9^ Despite this, the matrix and component chondrocytes of the ACC are often ignored or integrated into HAC/SCB analyses in OA research.^5,10^ This is likely due to the inability in most studies to reliably distinguish this ACC interface from the adjacent tissues. This persistent limitation in the field means that current interpretations of “subchondral changes” may in fact conflate two biologically distinct tissues. A critical need therefore exists to investigate the mechano-structural relationship between the ACC and SCB across the entire articular surface within the whole joint epiphysis.^11^

Historically, technical limitations have also hindered progress in our appreciation of the mechano-structural roles of ACC. Conventional histology, though widely used, is limited to two-dimensional insights and is subject to processing artefacts.^10^ Clinical and preclinical studies have primarily examined the thickness of articular cartilage and its association with lesion severity, yet reveal little about the internal sub-architecture that governs the biological integrity of the ACC. Consequently, there is a lack of research identifying microstructural determinants of the ACC/SCB osteochondral interface that may predispose the joint to functional compromise during OA development.^12^ Identification of such predisposing features could, however, have significant value as prognostic biomarkers.

Advanced imaging technologies now present an opportunity for transformative progress. High-resolution synchrotron computed tomography (sCT) enables unprecedented 3D visualisation of ACC microarchitecture.^11,13,14^ This, combined with digital volume correlation (DVC), extends the level of multiscale detail to capture the *entire* epiphyseal ACC compartment within intact loaded mouse joints and represents a significant methodological and conceptual advance, offering - for the first time - complete anatomical mapping of the ACC across entire condyles in healthy and OA joints. This will ultimately enable understanding of whether mechano-structural changes in OA could be primarily rooted within the ACC.

To interrogate these possibilities, we leveraged the predictable, spontaneous onset of knee OA in the medial tibial condyle of the murine STR/Ort strain, comparing outcomes to healthy ageing parental CBA controls.^15^ This model affords unique power to discriminate features that precede, accompany, and/or exacerbate OA pathological change in HAC. Using 3D whole-epiphysis, high-resolution sCT, we tested the hypothesis that structural features of tibial ACC differ fundamentally prior to knee OA onset, and that they may serve as predictive mechano-biomarkers or mechanistic drivers of OA progression in response to joint loading.

## Results

### Regionalised strain quantification by DVC reveals a selective unbalancing of compressive strain distributions across the STR/Ort ACC

To determine whether OA predisposition is associated with an anatomically biased joint response to physiological loading, regionalised DVC was conducted to quantify third principal (compressive) and first principal (tensile) strains within the medial and lateral ACC and SCB of STR/Ort and CBA tibial epiphyses. At 10-weeks of age, compressive strain magnitudes were greater in the lateral than medial ACC (Figure 1A,B; p=0.04) while no difference was evident in the SCB (Figure 2C) of STR/Ort mice. This latero-medial asymmetry was no longer evident in 40-week-old animals (Figure 1A, C-D). In contrast, compressive strains were evenly distributed across lateral and medial condyles within both the ACC and SCB of CBA tibial epiphyses at both ages (Figure 1B-E). A comparable lateral predominance was likewise observed for tensile strains in 10-week-old STR/Ort epiphyses (Figure 1F), with greater first principal strain magnitudes in the lateral ACC (Figure 1G) and SCB (Figure 1H, both p=0.04). In line with the measured compressive strains, tensile strains were also symmetrically distributed across the lateral and medial ACC and SCB in the epiphyses of 10- and 40-week-old CBA mice (Figure 1G-J).

**Figure 1:**
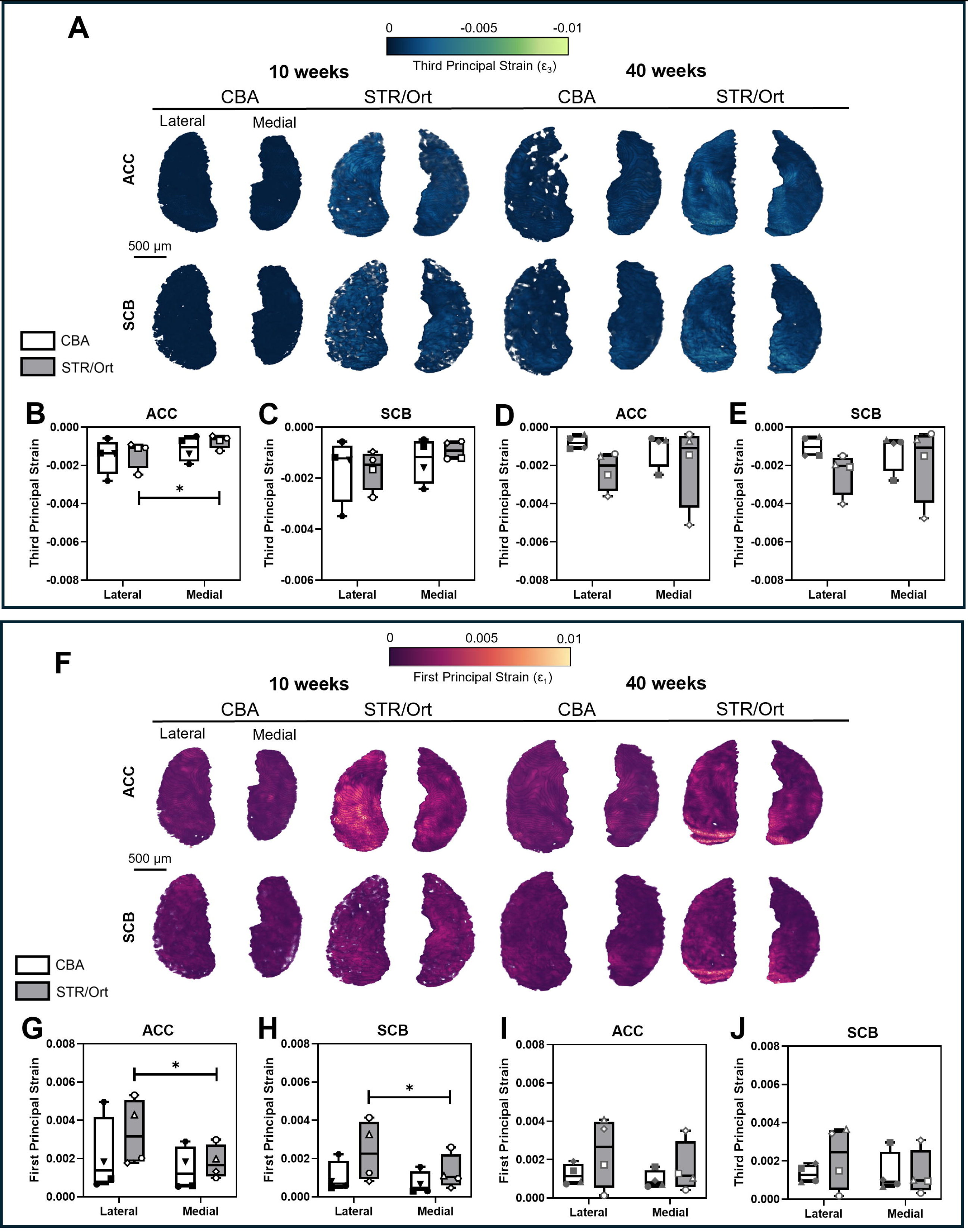
Regionalised strain quantification reveals the imbalanced distribution of compressive and tensile strain across the STR/Ort articular calcified cartilage (ACC) and subchondral bone (SCB). Compressive strains derived from DVC analyses are shown for the ACC (**A; top**) and SCB (**A; bottom**) from 10- and 40-week-old mice are presented as volume renders superimposed with strain magnitudes. Quantification of average compressive strain in the lateral and medial ACC and SCB of 10- (**B** and **C**, respectively) and 40-week-old (**D** and **E**, respectively) mice. Tensile strains generated in response to displacement-controlled loading within the ACC and SCB are shown (**F**), with average strain quantified in the lateral and medial ACC of 10- (**G** and **H**, respectively) and 40-week-old mice (I and J, respectively). Data are presented as box and whisker plots, where boxes represent the IQR, the central line denotes the median and the whiskers correspond to the minimum and maximum values. Symbols represent individual animals (N=4 per age and genotype). Statistical significance between condyles and genotypes was assessed using repeated measures two-way ANOVA with Šídák’s post hoc test (*p<0.05).

**Figure 2:**
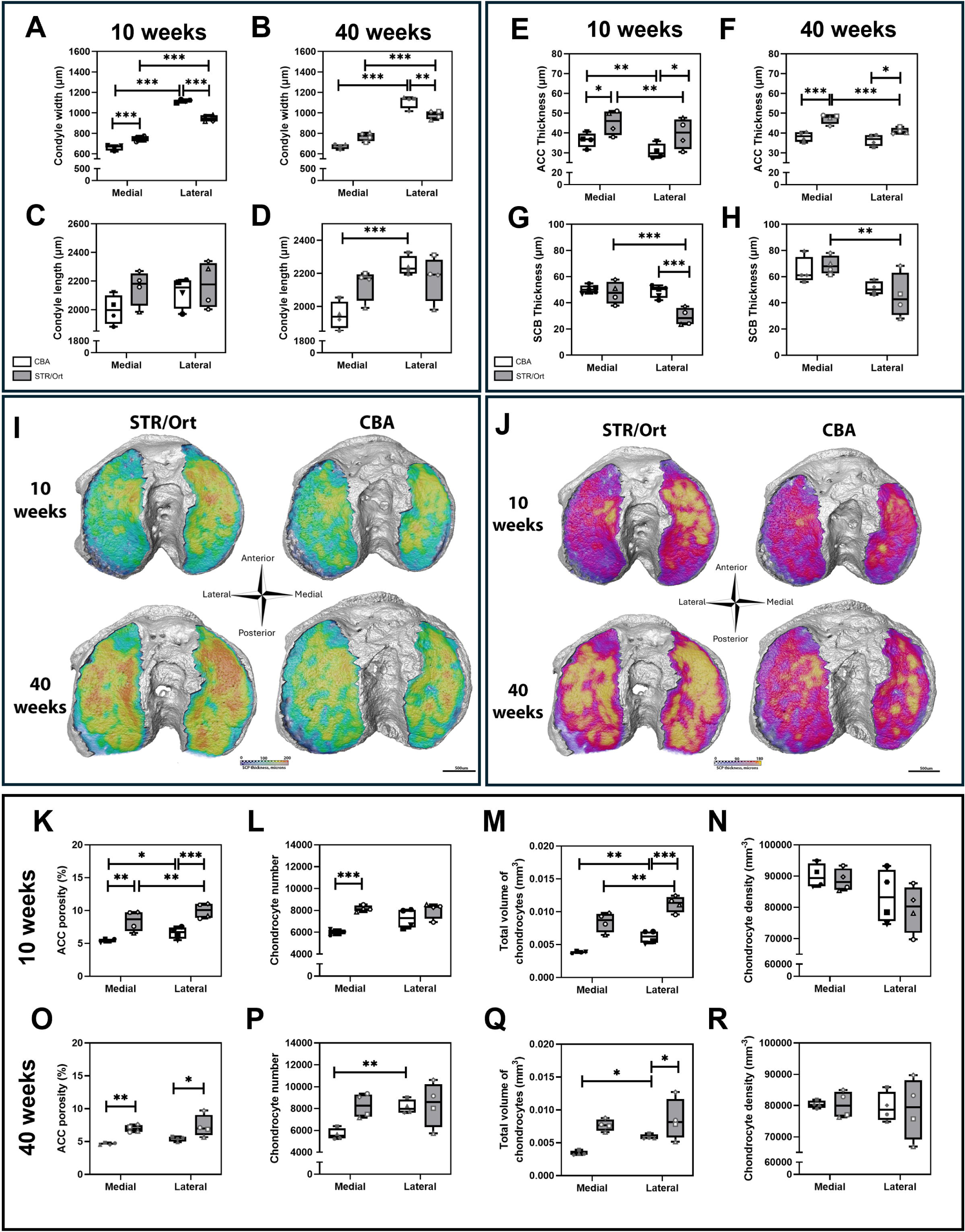
Gross anatomical evaluation of the articular calcified cartilage (ACC) and subchondral bone (SCB). Epiphyseal condylar width (µm) at **(A)** 10- weeks and **(B)** 40- weeks of age, and condylar length at **(C)** 10- weeks and **(D)** 40- weeks of age. Average thickness (µm) of the ACC **(E, F)** and SCB **(G, H)** at 10- and 40-weeks of age, respectively. 3D heatmaps of **(I)** ACC and **(J)** SCB thickness. Anatomical assessment of ACC porosity (%) at **(K)** 10-weeks and **(O)** 40-weeks of age, chondrocyte number at **(L)** 10-weeks and **(P)** 40-weeks of age, total volume of chondrocytes (mm^3^) at **(M)** 10-weeks and **(Q)** 40-weeks of age, and chondrocyte density (mm^-3^) at **(N)** 10-weeks and **(R)** 40-weeks of age. Data are presented as box and whisker plots, where boxes represent the IQR, the central line denotes the median and the whiskers correspond to the minimum and maximum values. Symbols represent individual animals (N=4 per age and genotype). Statistical significance between condyles and genotypes was assessed using two-way ANOVA with Fisher’s post hoc test (*p<0.05; **p<0.01; ***p<0.001).

Building on the compartment-specific strain patterns identified by DVC, regional FE modelling was used to determine whether these functional load-related differences were accompanied by changes in the predicted material properties of the ACC and SCB. In 10-week-old CBA epiphyses, stiffness was comparable between the lateral and medial condyles within both the ACC and SCB (Table 1). However, within each condyle, the SCB was significantly stiffer than the corresponding ACC (lateral, p=0.0003; medial, p<0.0001). A similar pattern was observed in 10-week-old STR/Ort epiphyses, in which ACC and SCB stiffness was comparable between the lateral and medial condyles, while the SCB was significantly stiffer than the overlying ACC in both condyles (both p=0.002).

**Table 1:** FE predictions of Young’s modulus and Poisson’s ratio for 10- and 40-week-old CBA and STR/Ort mice across the articular calcified cartilage (ACC) and subchondral bone (SCB) of the tibial epiphysis. Data presented as mean ± SD for N=3 mice per age and genotype. Statistical significance was assessed using two-way ANOVA with Šídák’s post hoc test. * denotes differences between CBA and STR/Ort mice, † denotes statistical significance between 10- and 40-week-old animals, and § denotes statistical significance between the ACC and SCB with single, double, triple and quadruple symbols corresponding to p<0.05, p<0.01, p<0.001 and p<0.0001, respectively.

|  |  |  | ACC |  | SCB |  |
| --- | --- | --- | --- | --- | --- | --- |
|  |  |  | Lateral | Medial | Lateral | Medial |
| Predicted Modulus<br>(GPa) | 10<br>weeks | CBA | $0.68 \pm 0.25$ | $0.58 \pm 0.03$ | $5.68 \pm 1.64$<br>§§§ | $6.52 \pm 0.67$<br>§§§§ |
| | | STR/Ort | $0.47 \pm 0.23$ | $0.42 \pm 0.15$ | $6.93 \pm 0.99$<br>§§ | $6.89 \pm 0.71$<br>§§ |
| | 40<br>weeks | CBA | $0.63 \pm 0.13$ | $0.37 \pm 0.06$<br>§ | $6.99 \pm 0.94$<br>§§ | $5.71 \pm 0.32$<br>§§ |
| | | STR/Ort | $0.11 \pm 0.02^{****},$<br>† | $0.09 \pm$<br>$0.04^{**}, \dagger$ | $1.95 \pm$<br>$1.35^{***}, \dagger\dagger\dagger$ | $1.4 \pm 0.97$<br>$^{**}, \dagger\dagger\dagger$ |
| Predicted Poisson's<br>Ratio (A.U.) | 10<br>weeks | CBA | $0.19 \pm 0.03$ | $0.19 \pm 0.02$ | $0.27 \pm 0.02$ | $0.23 \pm 0.01$ |
| | | STR/Ort | $0.21 \pm 0.05$ | $0.19 \pm 0.03$ | $0.28 \pm 0.03$ | $0.26 \pm 0.03$ |
| | 40<br>weeks | CBA | $0.18 \pm 0.02$ | $0.20 \pm 0.01$ | $0.25 \pm 0.05$ | $0.23 \pm$<br>$0.02$ |
| | | STR/Ort | $0.17 \pm 0.02$ | $0.24 \pm 0.04$ | $0.23 \pm 0.02$ | $0.26 \pm 0.02$ |

In 40-week-old CBA epiphyses, the ACC was stiffer in the lateral than the medial condyle (p=0.01), whereas the SCB stiffness remained comparable between condyles; the SCB also remained stiffer than the corresponding ACC in both lateral (p=0.0007) and medial (p=0.001) condyles. By contrast, 40-week-old STR/Ort epiphyses exhibited neither latero-medial differences in ACC or SCB moduli nor differences in stiffness between these compartments. At this advanced age, STR/Ort mice also exhibited lower predicted modulus than CBA mice across condyles and epiphyseal compartments (lateral ACC, p<0.0001, medial ACC, p=0.004, lateral SCB, p=0.0004 and medial SCB, p-0.001).

Ageing of CBA mice was accompanied by the preservation of ACC and SCB stiffness in both condyles, and this contrasted markedly with the clear age-related reduction in predicted modulus evident in both the lateral and medial ACC (p=0.01 and p<0.02, respectively) and SCB (p=0.0007 and p=0.0004, respectively) in STR/Ort mice. Compartmental differences between the ACC and SCB were not evident in 40-week-old STR/Ort. The predicted Poisson’s ratio remained comparable across genotypes, ages, condyles and epiphyseal compartments. Together, these findings indicate an age-associated loss of effective stiffness across both the ACC and SCB of OA-prone STR/Ort epiphyses, alongside an absence of compartment-specific stiffness differences in healthy joints.

### The ACC of OA-prone STR/Ort is thicker whilst the SCB is thinner than in healthy CBA mice

To understand the architectural relationships underpinning this imbalanced distribution of strains across the STR/Ort tibial osteochondral interface, we first assessed the gross condylar shape and composition of the ACC and SCB components. We found that the lateral condyle is significantly broader than the medial in both STR/Ort and CBA strains at 10- and 40-weeks of age (Figure 2A & B; p<0.001). However, comparison between mouse strains, showed that CBAs have a significantly broader lateral condyle than STR/Orts at both 10 (p<0.001) and at 40-weeks of age, Figure 2A & 2B). In contrast, the medial condyle in STR/Orts is significantly broader than in CBAs (p<0.01 at 10-weeks of age only, Figure 2A). No significant differences in condyle length were observed between strains at 10-weeks (Figure 2C), with asymmetry observed in CBA mice at 40-weeks of age (Figure 2D).

Detailed sub-segmentation of the subchondral plate (SCP) allowed quantification of the gross composition of its ACC and SCB components. This revealed that OA-prone STR/Ort mice have thicker ACC than healthy CBA mice at both 10-weeks (Figure 2E & I; p<0.05) and 40-weeks of age (Figure 2F & I; p<0.05). Condylar asymmetry was also observed in both strains at 10-weeks (Figure 2E; p<0.01), and this was only preserved in STR/Ort mice at 40-weeks of age (Figure 2F; p<0.01). In contrast, limited differences were observed in the SCB thickness, with STR/Ort mice showing asymmetry across condyles (Figure 2G & H; p<0.001) and strain differences only observed in the lateral condyles at 10-weeks of age (Figure 2G & J; p<0.001).

Overall, the thicker ACC coupled with a thinner SCB in STR/Ort mice, compared with CBA, indicates divergent SCP alterations that may have been obscured when these regions were analysed together. This therefore warranted the development of methods to further interrogate these more prominent shifts in the hitherto relatively undefined ACC.

### ACC in OA-prone STR/Ort mice contains chondrocyte lacunae that are morphologically distinct and more clustered than in healthily ageing joints

Microstructural evaluation of the ACC, enabled by sCT imaging of the entire joint revealed that STR/Ort have a more porous ACC than CBA mice across both condyles at 10- and 40- weeks of age (Figure 2K & O; p<0.05). Whilst a strain difference in chondrocyte lacunar number only existed on the medial condyle at 10-weeks of age (STR/Ort > CBA, Figure 3L; P<0.001), it was interesting to note that STR/ort mice appeared symmetrical in their condylar distribution of lacunae, whilst CBA possessed more in the lateral condyle in comparison to the medial (Figure 2L & P). These observations largely matched data on total chondrocyte lacunae volume (Figure 2M & Q), however no differences were found in chondrocyte lacunae density (Figure 2N & R). Together these data suggest that the ACC in STR/Ort contains more, larger-sized chondrocyte lacunae and thus greater ACC porosity than CBA mice.

**Figure 3:**
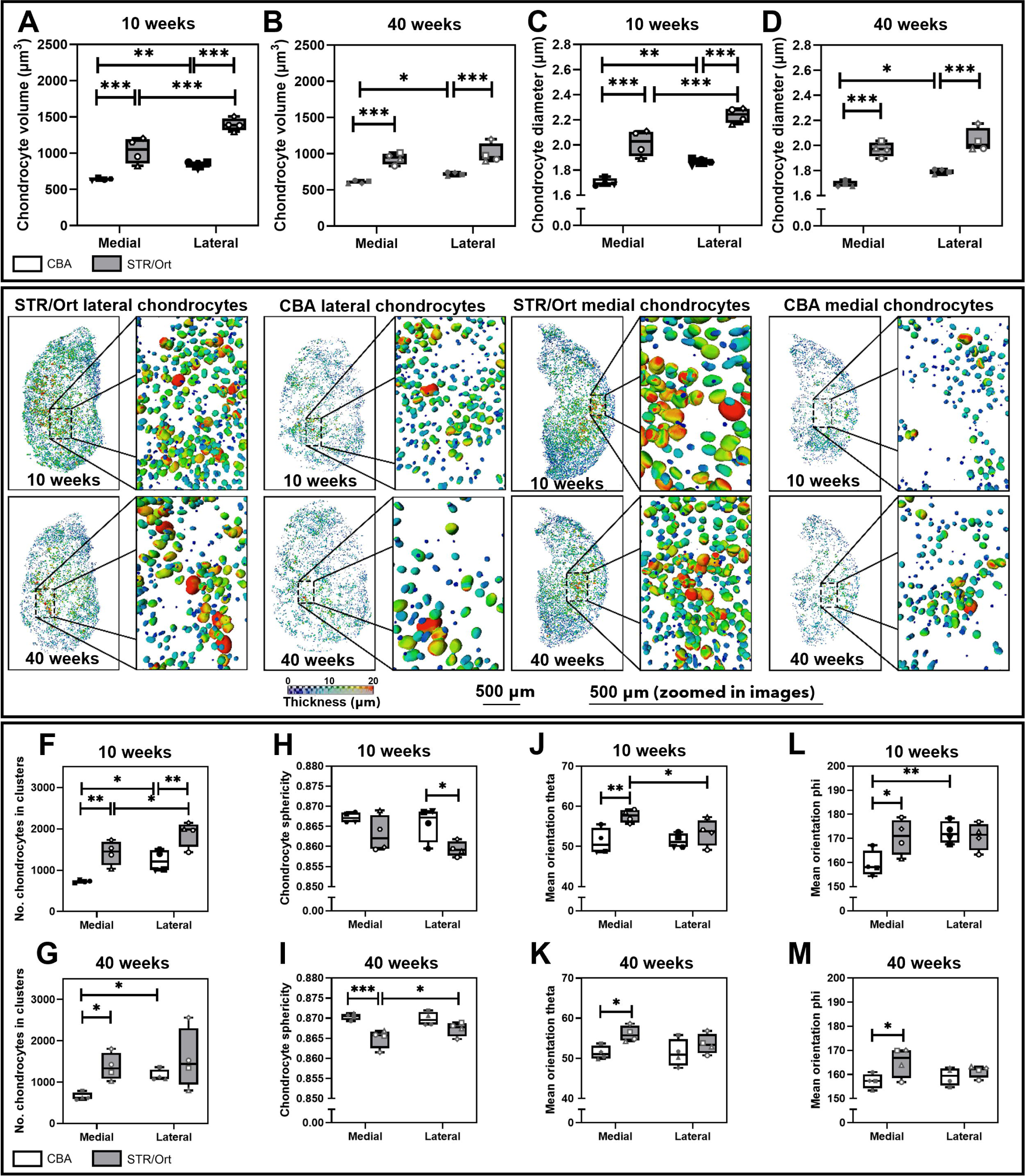
Microstructural analysis of the articular calcified cartilage in STR/Ort and CBA mice. Measures of chondrocyte volume (µm^3^) at **(A)** 10-weeks and **(B)** 40-weeks of age, and chondrocyte mean diameter (µm) at **(C)** 10-weeks and **(D)** 40-weeks of age. **(E)** 3D heatmap visualisations of the chondrocyte diameter from the superior surface to the inferior surface. Number of chondrocytes in clusters at **(F)** 10-weeks and **(G)** 40-weeks of age, chondrocyte sphericity at **(H)** 10-weeks and **(I)** 40-weeks of age, mean orientation theta at **(J)** 10-weeks and **(K)** 40-weeks of age, and mean orientation phi at **(L)** 10-weeks and **(M)** 40-weeks of age. Data are presented as box and whisker plots, where boxes represent the IQR, the central line denotes the median and the whiskers correspond to the minimum and maximum values. Symbols represent individual animals (N=4 per age and genotype). Statistical significance between condyles and genotypes was assessed using two-way ANOVA with Fisher’s post hoc test (*p<0.05; **p<0.01; ***p<0.001).

Indeed, closer interrogation of chondrocyte lacunar size revealed that STR/Ort ACC contains significantly larger chondrocyte lacunae than CBA mice, with measurements of greater mean lacunar volume (Figure 3A & B), and diameter (Figure 3C-E) at both ages. Compellingly, STR/Ort mice had significantly more clustered chondrocyte lacune than CBA comparators across both condyles at 10-weeks (Figure 3F, p<0.01), and specifically on the medial condyle at 40 weeks of age (Figure 3G, p<0.05). These observations were clear upon 3D reconstruction of ACC chondrocyte lacunae from within the medial and lateral condyles (Figure 3E). ACC lacunae in CBA mice were, by contrast, found to be more spherical, significantly so in the lateral condyle at 10 weeks of age (Figure 3H; p<0.05), and the medial at 40-weeks of age (Figure 3I; p<0.001). Further, analysis of chondrocyte orientation showed a significant increase in theta (Figure 3J & K) and phi (Figure 3L &M) in STR/Ort mice in comparison to CBA on the medial condyle at both ages, suggestive of chondrocytes tilting away from the vertical axis.

## Discussion

The mechano-structural relationship between the ACC and SCB remains unresolved and this has consequently limited our appreciation of whether these distinct tissues may serve as independent predictive mechano-biomarkers or mechanistic drivers of OA. STR/Ort mice are an extensively characterized preclinical model of age-related knee OA with multiple features comparable to those observed in primary human OA.^16^ Our recent research revealed that spatial microarchitectural SCP alterations preceded HAC loss and promoted focal subchondral strain concentration in STR/Ort joints.^17,18^ Here, we resolved full-field strains in the individual SCP components - ACC and SCB - and linked strain concentration to the underpinning microanatomy to question whether healthy joint function relies on the load-bearing properties and structural integrity of the ACC, and whether any specific ACC features are already present before OA onset, which may therefore be predictive of future pathology.^11,17^

Our findings demonstrate that ACC chondrocyte lacunae in the OA-prone STR/Ort were larger, more clustered, and morphologically distinct compared with those observed in healthy-ageing CBA controls, consistent with prior ex vivo findings linking hypertrophic, clustered ACC chondrocytes to an OA phenotype.^19–21^ As these differences were observed prior to OA onset in our model, they suggest that alterations in lacunar architecture are not merely consequences of degeneration but may reflect intrinsic differences in this strain, potentially linked to the known endochondral ossification defect detected in STR/Ort mice.^19^ In this context, a failure to adequately regulate lacunar size during growth may persist into adulthood. More broadly, these findings support the concept that OA may arise not simply through reactivation of embryonic endochondral pathways, but through their incomplete suppression or dysregulation following skeletal maturation.^20^ This interpretation reframes OA, at least in part, as a developmental disorder of cartilage maturation and mineralisation, with implications for how early-life tissue properties influence later disease risk.

The decreased lacunar sphericity observed herein is consistent with previous studies in human HAC.^22–24^ Our observation in the ACC of STR/Ort mice may indicate chondroptosis, followed by partial infilling of evacuated lacunae, providing a potential explanation for the age-related reduction in porosity and mean lacunar size observed particularly in the lateral condyle.^22,25^

Mechanical factors are well established as central to OA, yet the specific aspects of joint mechanics that initiate or drive disease onset and progression remain unclear.^26^ Our results suggest that functionally, lacunar morphology appears to have important biomechanical consequences for ACC behaviour. Excessively large or clustered lacunae may increase tissue compliance, leading to over-deformability under load. This, in turn, could compromise the ability of the ACC to protect the overlying HAC, particularly during repetitive joint loading, ultimately contributing to progressive tissue failure. Indeed, in our previous work, we revealed substantial differences in the mechanical response of healthy versus OA-predisposed murine ACC when subjected to mechanical loads.^11^ Our DVC findings here further highlight a mechanical dichotomy between healthy ageing and OA-prone murine joints, particularly in relation to the symmetry of load distribution. In OA-prone STR/Ort mice, strain patterns were less symmetric than in healthy CBA controls, suggesting misaligned load transfer across the joint. Given that joint loading is typically asymmetrical, differences in tibial condyle morphology between strains may further modulate the relationship between articular microstructure and disease. Indeed, the more symmetrical tibial plateau observed in STR/Ort, compared with the characteristically larger lateral condyle in CBA mice could predispose to mechanically unfavourable joint environments, potentially contributing to OA susceptibility, although causality remains to be established. While the present work focuses on structural descriptors, it is likely that underlying matrix composition, such as mineral density, collagen organisation, and non-collagenous protein content, also interacts with lacunar architecture to determine mechanical performance. These matrix constituents therefore represent important avenues for future investigation using methods such as Raman spectroscopy.^27,28^ Further, it would be interesting to examine strains around individual chondrocytes, particularly those regions with spatial localisation of hypertrophic chondrocytes.

It is also important to consider the limitations of our experimental and modelling approaches. Notably, soft tissues such as articular cartilage, menisci, ligaments, and other periarticular structures were not explicitly incorporated into the FE models. This omission reflects a technical constraint: DVC could not reliably resolve strains in these tissues due to insufficient phase contrast in the sCT imaging. However, it is well recognised that these soft tissues play critical roles in joint mechanics, particularly in the STR/Ort model, and their exclusion likely influences the interpretation of load transfer and strain distribution within the joint.^29^ A further limitation of our work is that this model represents a spontaneous, rather than injury-induced, form of age-related progressive OA, and thus whilst it shares important characteristics with human primary OA, it cannot be assumed that our findings will directly translate to human biology or anatomy and this warrants further imaging modality development. This is particularly given the highly heterogenous nature of OA in humans.^30^ Future studies could also investigate whether lacunar morphology can serve as an early biomarker of OA risk across models, including other primary OA systems such as the Dunkin–Hartley guinea pig or naturally aged C57BL/6 mice, to determine the generalisability and predictive value of these structural features.^31^ Further work could also consider the female STR/Ort mouse strain, as the STR/Ort mouse exhibits a sexually dimorphic OA phenotype by histology, with pathology consistently apparent in male mice, while females appear markedly protected.^32^

In conclusion, our work facilitates the “whole joint” organ approach by resolving regional nanoscale stains and tibial microarchitectures across the entire intact tibial epiphysis. Our data indicate that alterations in joint mechanics and tissue microstructure in the ACC are evident prior to OA onset. Specifically, we show that chondrocyte lacuna size and shape in the ACC is linked to asymmetrical strain distribution, defining these as mechano-predictive biomarkers of age-related OA.

## Methods

### Animals

All animal experiments were conducted in accordance with The UK Animals (Scientific Procedures) Act 1986 and were approved by the Institutional Animal Welfare Ethical Review Board at the Royal Veterinary College. Male STR/Ort mice were maintained as an in-house colony through brother-sister breeding and were examined at pre-OA stages in 8-11-week-old animals (N=4) and at advanced OA stages in 40-week-old animals (N=4).^15^ Age-matched male CBA mice (Charles River, UK) served as healthy parental-strain controls (N=4/age).^15^ All mice were housed in polypropylene cages under controlled environmental conditions of at 21 ± 2 °C and a 12 hr light/12 hr dark cycle and fed a standard RM1 maintenance diet (No.1; Special Diet Services, Witham, UK) and water ad libitum. Mice were euthanised by cervical dislocation, after which the left hindlimbs were dissected and the skin removed. Hindlimbs were wrapped in 1X phosphate buffered saline (PBS)-soaked gauze to maintain hydration prior to storage at -80°C until imaging.

### In situ loading

To conduct a full mechano-anatomical assessment of the ACC at the submicron level, we utilised our high-resolution in situ whole epiphyseal imaging and strain mapping framework in intact murine knee joints (Figure 4A).^17^

**Figure 4:**
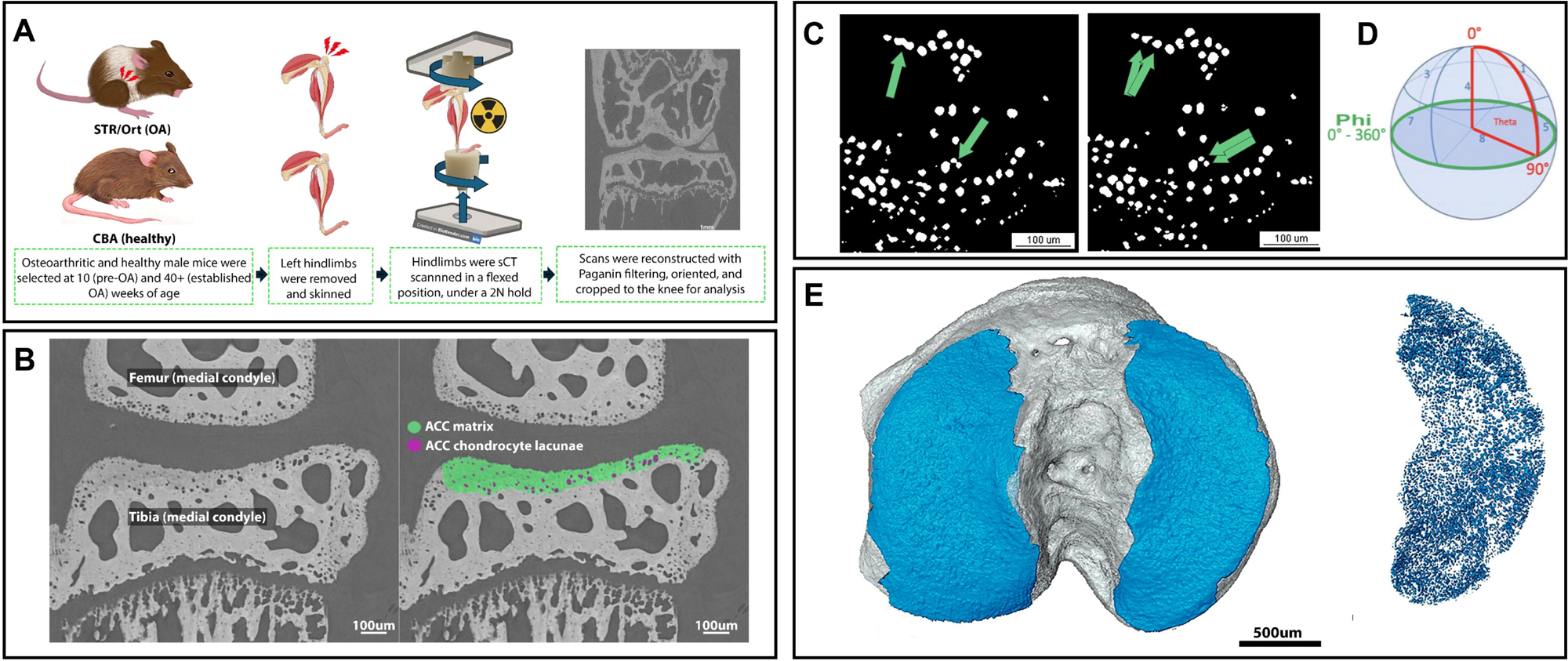
Characterization of ACC microarchitecture in the epiphyses of healthy CBA and osteoarthritis-prone STR/Ort knee joints with phase-contrast sCT. (A) Schematic of methodology representing isolation of hindlimbs from 10- and 40-week-old male CBA and STR/Ort mice and mounting in a Deben CT500 compression stage with a bespoke open frame, for in situ mechanical compression testing and sCT. (B) Segmentation of the ACC from sCT images. (C) Thresholding of resident chondrocytes with a watershed transformation to enable individual separation. (D) Calculation of chondrocyte orientation (Adapted from Bruker Method Note 145 with permission). (E) Example final volume renders of the ACC and chondrocytes for subsequent microarchitectural analysis.

In situ loading and synchrotron computed tomography (sCT) were performed according to a previously established protocol.^17^ Briefly, hindlimbs were thawed at room temperature and mounted in deep flexion between custom-made sample cups within a Deben CT500 in situ compression stage (Deben UK Ltd, UK), equipped with a bespoke open frame and a 100 N load cell (accuracy of ± 1%). Quasi dynamic axial compression was applied in the z-direction upwards from the lower cup prior to the acquisition of sCT images. A baseline pre-load of 2 N was applied to maintain deep flexion of the knee and ankle^16^ with the first series of sCT images acquired following a 15 minute stress relaxation period. Three further sCT images were acquired following incremental displacement of 20 µm, 70 µm and 170 µm, with each scan preceded by a 15-minute stress relaxation period (see Sharma et al., for loading curves).^17^

sCT datasets of in situ loaded joints were acquired using an isotropic voxel size of 1.45 µm on the BM05 beamline at the European Synchrotron Radiation Facility, Grenoble, France, (beamtime LS-3124) using a white X-ray beam filtered using 2.33 mm aluminium and 0.41 mm copper, producing an average energy of 63 keV.^17^ 6000 projections were collected in the half-acquisition fly-scan mode^33^ over 360 ° continuous rotation, using an angular step size of 0.06 ° with an exposure time of 18 milliseconds as previously described.^17^ sCT projections were flatfield and darkfield corrected prior to reconstruction using the in-house PyHST2 software package (version 2023a).^34^ Paganin phase-retrieval (δ/β=500)^35^ was applied prior to a 2D unsharp mask (σ=1, coefficient=3.5) in preparation for filtered back-projection and ring artefact removal.^36^ Finally, the reconstructed 32-bit volumes were converted to 16-bit TIFF stacks using a consistent grey-value range across all datasets.

### Digital Volume Correlation (DVC)

Reconstructed sCT volumes were cropped to encompass the knee joint in preparation for DVC as previously described.^17^ Briefly, the tibial epiphysis was segmented from the 2 N baseline load sCT dataset using a 3D region-growing algorithm in Avizo3D (version 2022.2, Thermo Fisher Scientific, USA). Osteocyte and chondrocyte lacunae filled using the “fill volume” function after which a 3-pixel dilation and erosion step was applied. The resulting binary segmentations were converted into simplified surfaces using a smoothing factor of 5 and subsequently used to generate unstructured finite element (FE) tetrahedral meshes.^34^ A nodal spacing of 30 voxels was selected based on optimisation using sequential repeat reference scans of a single knee joint acquired without further loading, as previously reported.^17^

The sCT datasets obtained following incremental displacements of 20 µm, 70 µm and 170 µm were rigidly registered to the corresponding 2 N baseline (reference) dataset for each knee joint. FE meshes were then generated for the registered reference and loaded image volumes and a global FE-based correlation was performed using the xDVC module in Avizo (version 2022.2, Thermo Fisher Scientific, USA). Upon convergence, displacement and strain fields were calculated at each mesh node, as previously described.^17^ Principal strain fields were visualised as volumetric maps using the scientific colour maps “Navia” for compressive strain and “Matter” for tensile strain.^37^

### ACC segmentation

Image stacks were opened in Avizo (version 2022.2, Thermo Fisher Scientific, USA). Following the contours of the cement line, ACC was manually segmented from SCB^38^ in the coronal plane in 36.25µm increments across the whole tibial plateau (Figure 4B). An interpolation operation was then applied to perform approximate segmentations of intervening regions. All interpolated regions were visually checked and any segmentation errors manually corrected. Care was taken to ensure that regions of non-articular calcified cartilage in close anatomical proximity (e.g. entheses) were excluded. A threshold was used to isolate the porous ACC mineral phase, and ‘fill slices’ operations were applied in coronal and sagittal planes to include pores (representing chondrocyte lacunae) in the final segmentation. ACC from medial and lateral condyles of each tibia were manually separated using 3D visualisation and manual segmentation.

Lacunae from each condyle were isolated for separate analysis via 3D thresholding, then subjected to a 2-pixel erode and dilate operation to remove any artefactual fragments of osteocyte lacunae at the cement line. These segmentations were then exported in .TIFF format, imported into CTAn software (Bruker, Belgium), subjected to a 2D shrinkwrap, saved as an image stack, then subjected to a watershed operation (tolerance 0) (Figure 4C). A single-pixel erode and dilate operation was applied to remove any remaining artefactual connections between distinct lacunae, and outcome images were saved as a “post-watershed” segmented lacunar image stack. Final segmentations were re-imported into Avizo for 3D visualisation to confirm that no significant artefacts were included. Chondrocyte orientation was calculated using 3D analysis in CTAn (Figure 4D). Together, this processing enabled volume renders of the ACC and chondrocyte lacunae for subsequent microarchitectural analysis (Figure 4E).

To isolate the strain fields generated within the ACC and SCB in response to in situ mechanical loading, volumetric bulk strain datasets were spatially registered and resampled to match the dimensions of the corresponding ACC and SCB segmentations. Interactive thresholding and masking were then applied to retain strain values within each anatomical compartment while excluding background voxels to generate compartment-specific strain volumes for subsequent analysis as previously described.^17^

### FE analysis of the ACC and SCB

FE models were generated from sCT-derived anatomical segmentations of the ACC and SCB for each CBA and STR/Ort knee joint using the procedure described previously.^17^ Briefly, segmentations of the ACC and SCB from individual CBA and STR/Ort tibial epiphyses were converted into surfaces in Avizo3D and discretised into unstructured tetrahedral meshes with nodal spacing of 25 voxels. Homogenous mechanical properties of elastic modulus of 150 MPa and Poisson’s ratio of 0.3 were used. Effective material properties were estimated using an inverse optimisation approach implemented in MATLAB by minimising the root-mean-square error between spatially corresponding FE-predicted and DVC-derived principal strain fields.^39^ Model geometry, boundary conditions and applied displacements were held constant throughout optimisation. Full details of mesh generation, model implementation and material property optimisation are provided in Sharma et al., 2026.^17^

### Statistical analysis

All investigators were blinded to the genotype and age of the animals until data analysis. Normality was assessed using the Shapiro-Wilk test. Statistical analyses were performed in GraphPad Prism (version 10.5.0; Massachusetts, USA). Post-watershed lacunar segmentations were used for all lacunae analyses, with pre-watershed segmentations used only as a comparator to analyse chondrocyte clustering. Two-way ANOVA was conducted to determine the independent and interactive effect of genotype and age on strain within the ACC and SCB with post hoc tests conducted for further comparisons.

## Data availability

The processed sCT datasets, DVC-derived strain fields, and FE models are available from the corresponding authors upon reasonable request.

## Acknowledgements

Laboratory facilities were provided by the Royal Veterinary College, the University of Brighton and the Research Complex at Harwell. We acknowledge the European Synchrotron Radiation Facility for provision of synchrotron radiation facilities under proposal number LS-3124. We thank C. Reinhardt and L. Sinclair (University of Manchester at Harwell) for access to the Deben CT500 mechanical testing rig and E. Burke O’Leary, C. Berruyer and V. Schoeppler for their assistance during beamtime. We thank R. Chang, P. Salmon and J. Trend for their discussions about the analysis of the data. We thank the Histology SRF at the University of Liverpool for their technical assistance.

This work was supported by the UKRI MRC (MR/V033506/1), the BLAST Network (BB/W01825X/1), Royal Academy of Engineering (CiET 1819/10) and the Chan Zuckerberg Initiative (CZIF2021-006424 and CZIF2022-316777). For the purpose of open access, the authors have applied a Creative Commons Attribution (CC-BY) license to any Author Accepted Manuscript version arising from this submission.

## Author contributions

Conceptualisation: LAEE, AS, PDL, AAP, KAS; Methodology: LAEE, AS, LEB, KM, JC, ALP, PDL, AAP, KAS; Software: LAEE, AS, LEB, KM, JC, JB; Validation: LAEE, AS; Formal analysis: LAEE, AS, LEB, JC; Investigation: LAEE, AS, LEB, JC, ALP, JB; Resources: PDL, AAP and KAS; Data curation: LAEE, AS; Writing – original draft: LAEE, AS, KAS; Writing – review and editing: all authors; Visualisation: LAEE, AS, JC; Supervision: PDL, AAP and KAS; Project administration: PDL, AAP and KAS; Funding acquisition: PDL, AAP and KAS.

